# Hypoxia-activated *scleraxis a* mediates epicardial progenitor differentiation into a unique cardiac perivascular cell type

**DOI:** 10.64898/2026.02.26.708296

**Authors:** Björn Perder, Yu Xia, Jun Yao, Miaoyan Qiu, Alvin Gea Chen Yao, Muhammad Naeem, Paul Zumbo, Ignace Van der Wee, Avi Yakubov, Kazu Kikuchi, Doron Betel, Todd Evans, Michael R. Harrison, Jingli Cao

## Abstract

The epicardium is a crucial source of progenitor cells and paracrine signals that support heart development and regeneration. However, the molecular mechanisms that guide epicardial cell fate decisions remain incompletely understood. Here, we identify the transcription factor Scleraxis a (encoded by *scxa*) as a key regulator of epicardial progenitor differentiation in zebrafish. Through single-cell transcriptomics, genetic lineage tracing, and cardiac injury models, we show that *scxa* is transiently induced in activated epicardial progenitor cells (aEPCs) during both heart regeneration and developmental coronary angiogenesis. *scxa*^+^ epicardial cells primarily give rise to a previously uncharacterized cardiac population of perivascular cells marked by *col18a1a*, molecularly distinct from classical pericytes and vascular smooth muscle cells. We refer to this population as epicardial-derived perivascular mesenchymal cells (Epi-PMCs). These Epi-PMCs closely associate with coronary vessels and may contribute to vascular stabilization and remodeling, potentially through the anti-angiogenic but vessel-stabilizing activity of endostatin derived from collagen XVIII. Loss of *scxa* increases coronary vessel density. Mechanistically, we identify hypoxia and Hif1a signaling as upstream regulators of *scxa*, with systemic hypoxia or Hif factor stabilization robustly inducing *scxa* expression in the epicardium. Together, these findings uncover a hypoxia-responsive Scxa-Col18a1a axis that drives epicardial differentiation toward a vascular-supportive fate, offering new insight into the regulation of coronary vessel development and the regenerative potential of the epicardium.

## INTRODUCTION

The epicardium, a thin layer of mesothelial cells surrounding the heart, plays a pivotal role in heart development and regeneration. During embryonic heart development, the epicardium gives rise to various cell types, including fibroblasts, vascular smooth muscle cells (VSMCs), and pericytes, making it a critical source of progenitors essential for coronary vessel formation and myocardial development^1,2^. Several signaling pathways and factors have been implicated in regulating epicardial cell differentiation. For instance, TGF-β promotes differentiation into fibroblasts and VSMCs, while Wnt or Notch signaling drives the VSMC fate^3–6^. PDGF signaling was reported to guide epicardial-to-mesenchymal transition (EMT) and differentiation into VSMCs and pericytes^7,8^. Transcription factors like Tcf21 and WT1 are crucial for epicardial progenitor specification, EMT, and fibroblast lineage commitment^9–12^. In response to cardiac injury, the epicardium is reactivated, reinitiating a developmental program that supports tissue repair and regeneration by contributing cells and releasing paracrine factors^2,13,14^. During regeneration, several subpopulations of epicardial cells emerge including an activated epicardial progenitor cell type (aEPC) that we recently identified^15^. In addition to renewing the epithelial epicardium, these aEPCs migrate to enclose the wound, undergo epithelial-mesenchymal transition (EMT), and differentiate into *pdgfrb*^+^ mural cells and *hapln1a*^+^*pdgfra*^+^ mesenchymal cells^15^. aEPCs upregulate a specific transcriptional program upon heart injury, including extracellular matrix (ECM) component gene *col12a1b* and pentraxin-related protein 3 (*ptx3a*)^15^. Despite these insights, it remains elusive what molecular and environmental cues determine whether epicardial progenitors differentiate into fibroblasts, VSMCs, pericytes, or other cell types and how epicardial cells coordinate with endothelial cells to form coronary vessels. Addressing these questions is critical for clarifying the regulatory mechanisms of epicardial differentiation and could significantly enhance strategies for tissue engineering and regenerative medicine.

Scleraxis (Scx) is a basic helix-loop-helix (bHLH) transcription factor best known for its pivotal role in embryonic mesodermal development, particularly in promoting tendon formation and tendon cell differentiation^16–20^. Beyond its established role as a marker of tendon cell fate, Scx has broader functions in regulating cellular differentiation and tissue remodeling. In the heart, Scx was identified in the developing cardiac valves in mice, where it plays a key role in cell differentiation and extracellular matrix remodeling during valvulogenesis^21,22^. Scx expression has also been observed in the pro-epicardial organ and embryonic epicardial cells and is downregulated after birth^23–25^. *Scx* knockout mice exhibit ∼50% fewer cardiac fibroblasts, suggesting a role in fibroblast lineage commitment^26^, yet another study propose an endothelial fate for Scx^+^ pro-epicardial cells^23^, underscoring the need to clarify its diverse functions in epicardial and cardiac development. In addition, loss of *Scx* in mice results in hypocellular hearts, while prolonged post-injury *Scx* activation drives fibrotic myofibroblast differentiation^26^. These context-dependent effects highlight that Scx functions are shaped by local signaling and lineage states.

In the current study, we investigated the role of zebrafish scleraxis a (*scxa*) in aEPC cell fate and its contribution to heart development and regeneration. Using single-cell transcriptomics, we identified *scxa* expression in aEPCs and their mesenchymal progenies, with transient expression during coronary angiogenesis and strong induction under hypoxic conditions. Lineage tracing demonstrated that ventricular *scxa*^+^ cells originate from epicardial progenitors and mainly differentiate into a previously uncharacterized *col18a1a*^+^ perivascular cell population distinct from pericytes and VSMCs that we refer to as epicardial-derived perivascular mesenchymal cells (Epi-PMCs). Our findings reveal that hypoxia-driven *scxa* expression supports epicardial-to-perivascular differentiation, supporting coronary vasculature formation. We thus uncover a new transcriptional regulatory program for epicardial cell differentiation.

## RESULTS

### *scxa* is expressed in differentiating epicardial progenitors in heart regeneration

In a search for regulators of epicardial cell differentiation, we looked for transcription factors enriched in aEPCs in our single-cell transcriptomic data^15^. We found that the transcription factor *scxa* is expressed in a portion of the *ptx3a*^+^*col12a1b*^+^ aEPC cluster, as well as some epicardial progeny that take on a mesenchymal cell fate (*hapln1a*^+^) after heart injury (Fig. 1a, 1b, and Fig. S1). *scxa* expression partially overlaps with *col12a1b* (Fig. 1c). To confirm *scxa* expression, we crossed a *scxa:mCherry* BAC (bacteria artificial chromosome) reporter^27^ with the *tcf21:nucEGFP* line, which labels both the quiescent and activated epicardial cells and their progenies^15^. Without an injury, no or minimal *scxa*:mCherry signal was detected on the ventricular surface of adult hearts. By contrast, we observed prominent *scxa* expression at 3 days post-ventricular amputation injury (dpa) in the boundary zone of the wound, colocalizing with *tcf21*:nucEGFP (Fig. 1d). Co-expression of *scxa* and *tcf21* is further confirmed in cryosections of hearts at 3 dpa. Notably, many of these *scxa*^+^ *tcf21*^+^ cells reside in the compacted myocardium, indicating a mesenchymal, rather than epithelial, identity (Fig. 1e).

**Figure 1.**
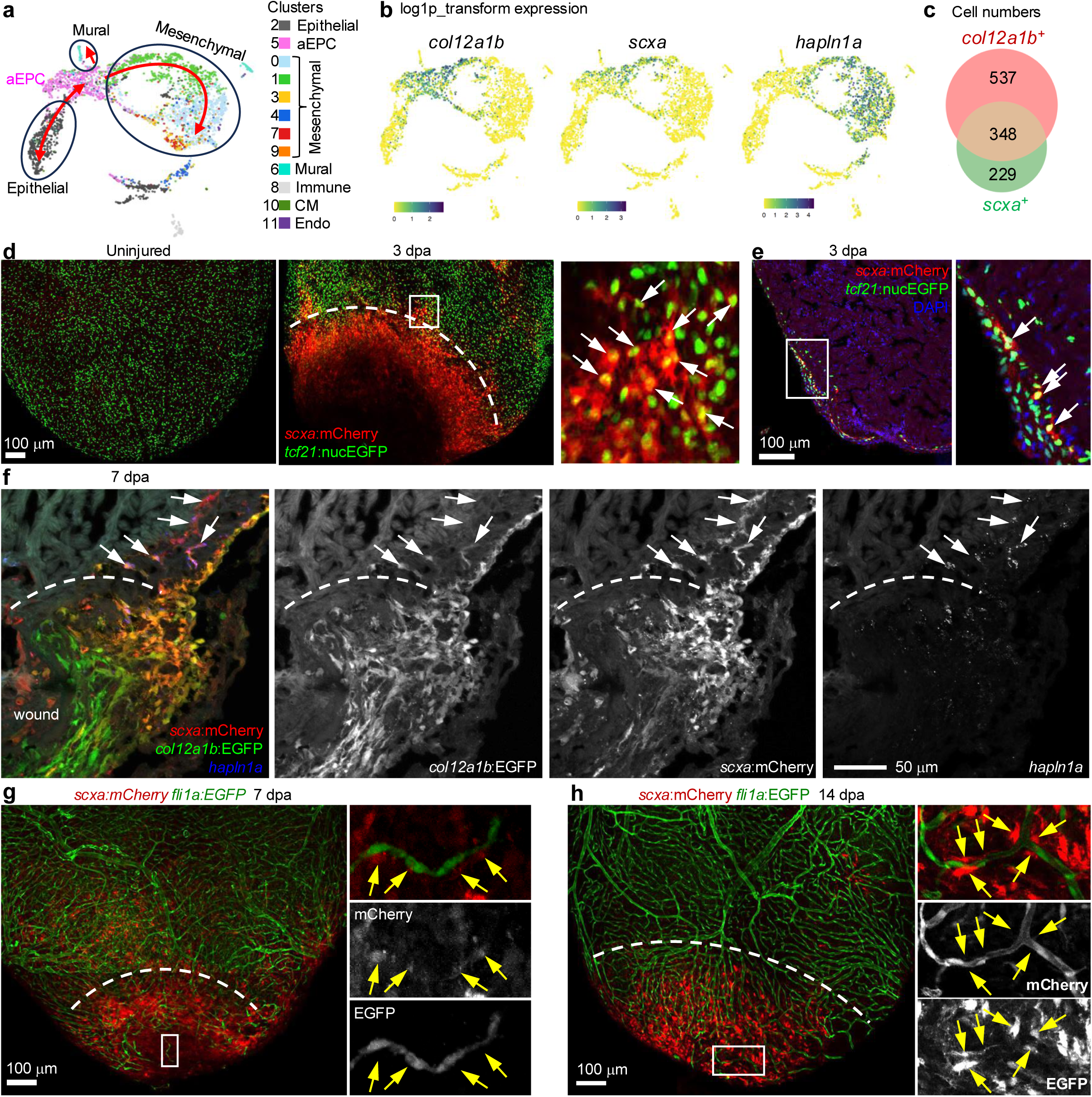
*scxa* is expressed in differentiating epicardial progenitors during heart regeneration. **a**, Cell trajectories suggested by pseudotime analysis with Monocle 3 is shown on a UMAP. **b**, Normalized expression (log1p transformed expression) of selected genes on UMAPs. **c**, A Venn diagram showing numbers of cells expressing *col12a1b* and/or *scxa* at 3 dpa. **d**, Whole-mount images of the ventricular surface showing expression of *tcf21*:nucEGFP (green) and *scxa*:mCherry (red) at 3 dpa. The framed region is enlarged to show details on the right. Arrows indicate representative mCherry^+^ EGFP^+^ cells. Scale bar, 100 μm. **e**, A section image of a 3 dpa heart showing *tcf21*:nucEGFP (green) and *scxa*:mCherry (red) expression. DAPI staining is shown in blue. The framed region is enlarged to show details on the right. Arrows denote representative mCherry^+^ EGFP^+^ cells. Scale bar, 100 μm. **f**, A section image of a 7 dpa heart showing *col12a1b*:EGFP (green) and *scxa*:mCherry (red) expression. HCR staining of *hapln1a* is shown in blue. Single-channel images are shown in grayscale. Arrows denote representative mCherry^+^ *hapln1a*^+^ cells. Scale bar, 50 μm. **g**, **h**, Whole-mount images of the ventricular surface showing expression of *fli1a*:EGFP (green) and *scxa*:mCherry (red) at 7 dpa (g) and 14 (h). The framed regions are enlarged to show details on the right with single-channel images shown in grayscale. Arrows indicate representative mCherry^+^ cells adjacent to vessels. Scale bars, 100 μm.

We previously identified *col12a1b* as a marker for aEPCs in zebrafish heart regeneration (Fig. S1)^15^. To confirm *scxa* expression in aEPCs, we crossed the *scxa:mCherry* line with the aEPC reporter *col12a1b:EGFP*. As shown in Fig. 1f, *scxa* expression partially overlapped with that of the aEPC marker *col12a1b* at the boundary zone of the wound at 7 dpa, while *scxa*^-^*col12a1b*^+^ aEPCs occupied the wound. We wondered whether some *scxa*^+^ cells gain mesenchymal identity by expressing the mesenchymal epicardial cell marker *hapln1a*^15^. We thus performed hybridization chain reaction (HCR) staining targeting *hapln1a* on cryosection samples from 7 dpa. As expected, some of *scxa*^+^ cells in the inner layer of the ventricular wall turned off *col12a1b* expression but gained *hapln1a* expression (Fig. 1f). These expression patterns perfectly recapitulate our scRNA-seq result confirming that *scxa* transcripts are found both in the *col12a1b*^+^ aEPCs as well as their mesenchymal progeny. This suggests that acquisition of *scxa* expression in aEPCs accompanies their transition to a mesenchymal cell fate.

The *hapln1a*^+^ mesenchymal epicardial cells are reported to support compact myocardium and coronary vessel growth and regeneration^15,28,29^. To characterize the *scxa*^+^ mesenchymal epicardial cells, we analyzed the *scxa* expression in hearts carrying the pan-endothelial reporter *fli1a:EGFP* to visualize revascularization. We observed that many *scxa*^+^ cells are associated with regrowing blood vessel sprouts in the wound site at 7 dpa (Fig. 1g). By 14 dpa, heart wounds were still enriched in *scxa*^+^ cells where they form a network surrounding the regenerating vasculature (Fig. 1h). *scxa*^+^ cells are seen frequently contacting coronary blood vessels (inserts in Fig. 1h), indicating a perivascular identity. Taken together, these results suggest that *scxa* is associated with epicardial cell differentiation to perivascular cells during heart regeneration.

### Transient *scxa*^+^ epicardial cells are associated with coronary angiogenesis in heart development

The previously unperceived vascular association of *scxa*^+^ epicardial lineage is intriguing. To better evaluate the relationship between *scxa*-expressing epicardial cells and the coronary vasculature, independent of the inflammation and wound healing processes observed during heart regeneration, we next characterized the spatiotemporal expression pattern of *scxa* during coronary angiogenesis, which begins in the juvenile stage at around 5-6 weeks post fertilization (wpf)^30^. To determine whether *scxa*^+^ cells exhibit characteristics of activated epicardial progenitor cells (aEPCs), we analyzed their co-expression with the aEPC marker *col12a1b*. Before 4 wpf, ventricular *scxa* expression and *col12a1b* expression were undetectable. At approximately 4 wpf, *col12a1b*:EGFP expression emerged in epicardial cells prior to the onset of *scxa*:mCherry expression, predominantly near the atrioventricular canal (AVC), where coronary vessel sprouts are known to initiate (Fig. 2a) ^30^. However, coronary vessels had not developed at this stage. At around 5 wpf, ventricular *scxa* expression was observed in *col12a1b*^+^ cells near the AVC (Fig. 2b). This suggests that *col12a1b* activation in aEPCs precedes *scxa* expression, mirroring the pseudotime trajectory observed in heart regeneration.

**Figure 2.**
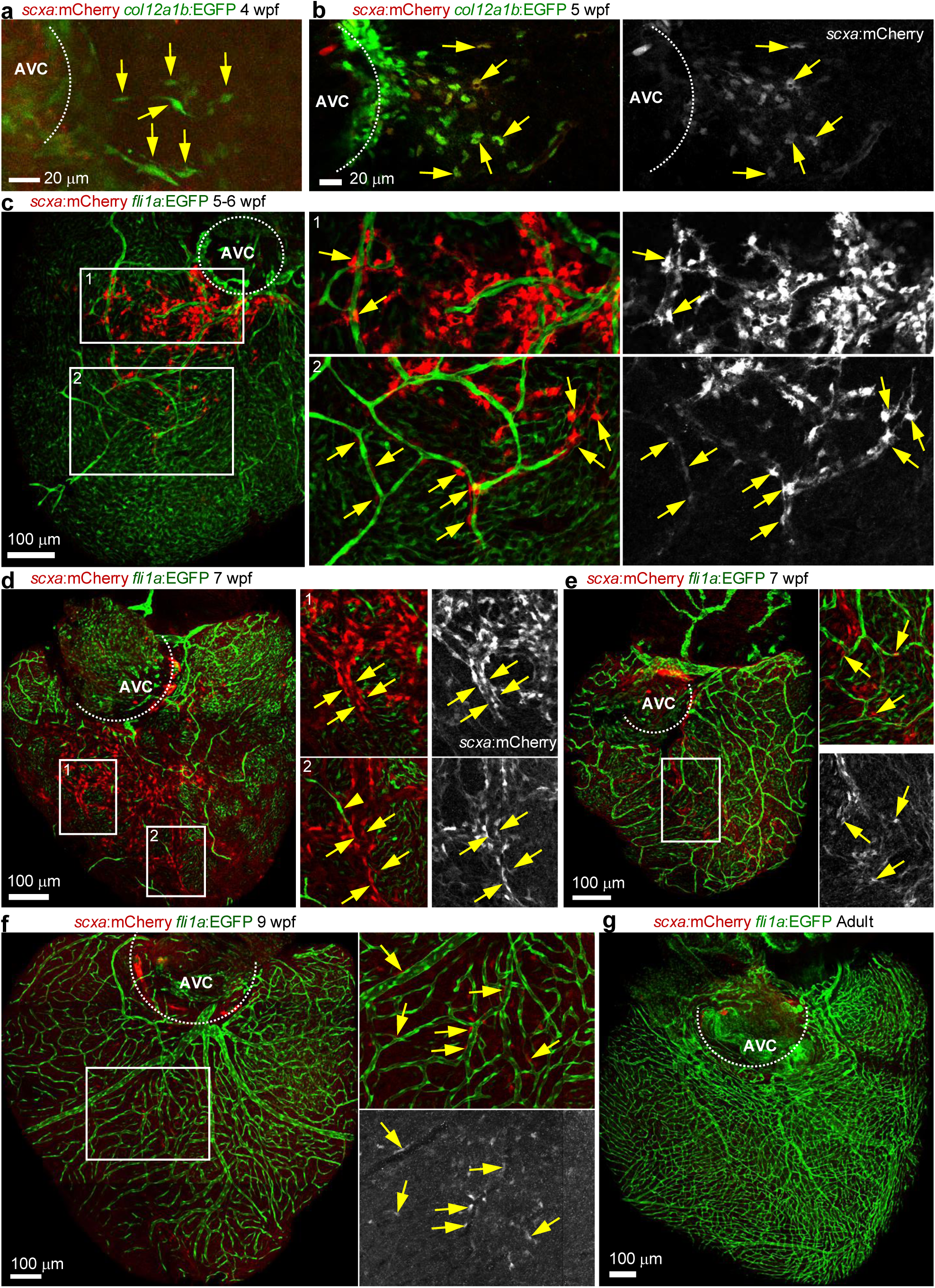
Transient *scxa*^+^ epicardial cells are associated with coronary angiogenesis. **a**, **b**, Whole-mount images showing *scxa*:mCherry and *col12a1b*:EGFP expression at 4 wpf (a) and 5 wpf (b). Arrows indicate representative cells. AVC, atrioventricular canal. Dashed lines indicate approximate AVC regions. **c,** Whole-mount images showing *scxa*:mCherry (red) and *fli1a*:EGFP (green) expression at 5-6 wpf, with framed regions enlarged on the right. Single channel images are shown in grayscale. The dashed line indicates approximate AVC. Arrows denote representative *scxa*^+^ cells. Single-channel images are shown in grayscale. Scale bars, 100 μm. **d**, **e,** Whole-mount images showing *scxa*:mCherry (red) and *fli1a*:EGFP (green) expression at around 7 wpf, with framed regions enlarged on the right. Single-channel images are shown in grayscale. Dashed lines indicate approximate AVC regions. Arrows mark *scxa*^+^ cells, while the arrowhead indicate an endothelial tip cell. Scale bars, 100 μm. **f,** Whole-mount images showing *scxa*:mCherry (red) and *fli1a*:EGFP (green) expression at around 9 wpf, with framed regions enlarged on the right. The single channel image is shown in grayscale. The dashed line indicates approximate AVC. Arrows denote representative *scxa*^+^ cells. Scale bars, 100 μm. **g**, Whole-mount images showing *scxa*:mCherry (red) and *fli1a*:EGFP (green) expression in the adult heart. The dashed line indicates approximate AVC. Scale bars, 100 μm.

At 5-6 wpf, all hearts exhibiting *scxa* expression contained a subset of *scxa*^+^ cells associated with the vasculature (Fig. 2c). Vessel-associated *scxa*^+^ cells were observed either in direct contact with the vessel or as part of a surrounding cellular ensemble (insets, Fig. 2c). Interestingly, in ventricular regions not yet vascularized, clusters of *scxa*^+^ cells were organized in parallel orientations. Endothelial tips from nearby vasculature appeared to orient toward these *scxa*^+^ clusters, suggesting directed growth of vessel sprouts toward these cells (Fig. 2c, frame 2 right side). This expression pattern continued to 7 wpf (Fig. 2d, frames) when *scxa* expression peaked. Decreased *scxa* expression was observed along with increased coronary vessel coverage from 7 to 9 wpf (Fig. 2e, f). In adult hearts with an established coronary vasculature, ventricular *scxa* expression was minimal (Fig. 2g). These observations suggest that transient *scxa^+^* epicardial cells, present before the emergence of endothelial sprouts and during vessel growth, may facilitate coronary angiogenesis.

### *scxa*^+^ epicardial cells primarily give rise to *col18a1a*^+^ mesenchymal cells

Noting the correlation of *scxa*^+^ cells with angiogenesis and their transient presence, we sought to determine their identity and progenies. We first explored the origin of ventricular *scxa*^+^ cells, and trajectory analysis indicated that these cells arise from the epithelial layer of the epicardium (Fig. 3a). To test this, we performed lineage tracing using animals carrying the *tcf21:CreER^t2^*, *ubb:loxP-TagBFP-stop-loxP-EGFP*, and *scxa:mCherry* transgenes^31,32^ (Fig. 3b). Following 4-hydroxytamoxifen (4HT) treatment during embryonic stages (2-4 dpf), analysis at 5 wpf revealed that all ventricular *scxa*^+^ cells expressed Cre-induced EGFP^+^, indicating their derivation from epicardial or pro-epicardial cells. We previously demonstrated that epicardial progenitor cells (aEPCs) give rise to *pdgfrb*^+^ pericytes or VSMCs during heart regeneration (Fig. 3a)^15^. Given the pericyte-like morphology of epicardial cell derived *scxa*^+^ perivascular cells, we crossed *scxa:mCherry* fish with the *pdgfrb:EGFP* line that labels pericytes and VSMCs^33^. Unlike pericytes that express high levels of *pdgfrb* and feature a characteristic elongated morphology, *scxa*^+^ perivascular cells mostly had a much lower-level expression of *pdgfrb* and exhibited a relatively round morphology with cell protrusions (Fig. 3c, arrows). These differences suggest that *scxa* is expressed in a perivascular cell population distinct from pericytes or VSMCs. To define their identity, we searched for new markers in our scRNA-seq data. We found that *col18a1a* is highly enriched in the mesenchymal progenies of aEPCs but not in pericytes and vSMCs (Fig. 3a and Fig. S1). COL18A1 is primarily localized in the basement membrane of blood vessels and epithelial tissues to maintain the structural integrity^34,35^. COL18A1 can be cleaved into endostatin peptide, which inhibits angiogenesis and stabilizes newly formed endothelial tubes^36–38^. However, the cellular source and function of COL18A1 in coronary vasculature development remains unclear. To confirm whether *scxa*^+^ cells give rise to *col18a1a*^+^ cells in the heart ventricle, we performed lineage-tracing employing an inducible *scxa:CreER^t2^* line (Ref. ^39^) crossed with the *ubb:loxP-TagBFP-stop-loxP-EGFP* reporter and applied labeling at juvenile stages with 3 overnight 4HT treatments at 6, 7 and 8 wpf (Fig. 3d). HCR staining of adult hearts confirmed that most (∼72%) of the lineage-traced progenies express *col18a1a* with low or no *pdgfrb* expression, which matched our prediction (Fig. 3a, e, g). We observed ∼14% of EGFP cell expressing high level of *pdgrb*^high^ indicating a mural cell identity (Fig. 3e, frame 2 and Fig. 3g). *col18a1a* expression was absent in *pdgfrb*^high^ mural cells (Fig. 3e, frame 1), further indicating that *col18a1a*^+^ cells are distinct from mural cells. In addition, we also observed ∼14% of EGFP^+^*col18a1a*^-^*pdgfrb*^-^ progenies (Fig. 3g), likely other mesenchymal progeny or *scxa*^+^ cells yet to differentiate. These observations indicate that *col18a1a*^+^ cells represent the dominant fate of *scxa*^+^ aEPCs, with mural cells as a minor fate. Moreover, *scxa^+^*and *col18a1a*^+^ cells also express *vegfaa* (Fig. 3a, f), suggesting a potential pro-angiogenic function of these cells. We next examined the fate of *scxa^+^*cells following heart injury. Adult fish carrying the *scxa:CreER^t2^*; *ubb:loxP-TagBFP-stop-loxP-EGFP* transgenes were subjected to ventricular amputation, followed by 4HT treatment at 6-8 dpa, when *scxa* expression is elevated in the wound region. Hearts were collected at 14 dpa (Fig. 3h). We found that approximately 57% of EGFP^+^ cells expressed *col18a1a* with low or no *pdgfrb* expression, consistent with an Epi-PMC identify (Fig. 3i, arrows; Fig. 3j). About 1.7% of EGFP^+^ cell are *col18a1a^-^ pdgrb*^high^ mural cells, and the remaining ∼41% were *col18a1a*^-^*pdgfrb*^-^ (Fig. 3i, j). In conclusion, these findings suggest that transient *scxa* expression in the epicardium is primarily associated with epicardial differentiation towards a *col18a1a^+^* cell fate, which may support coronary vasculature formation.

**Figure 3.**
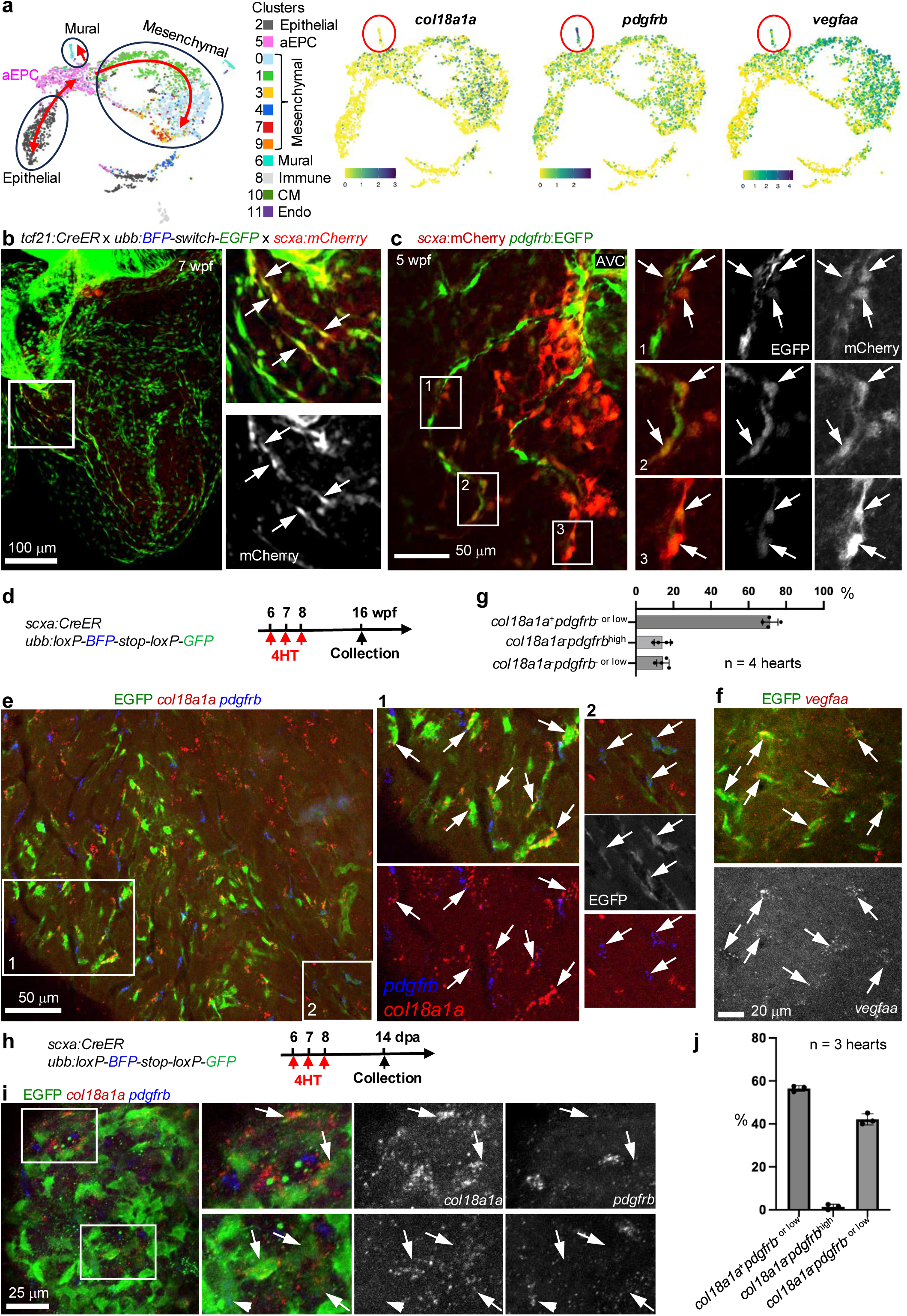
*scxa*^+^ epicardial cells give rise to *col18a1a*^+^ cells. **a**, Cell trajectories suggested by pseudotime analysis with Monocle 3 is shown on a UMAP. Normalized expression (log1p transformed expression) of selected genes (*col18a1a*, *vegfaa*, and *pdgfrb*) are on UMAPs, with the mural cell cluster circled. **b,** Whole-mount images showing *scxa*:mCherry (red) and epicardial progenies (EGFP^+^, green) at 7 wpf. Scale bars, 20 μm. **c,** Whole-mount images showing *scxa*:mCherry (red) and *pdgfrb*:EGFP (green) expression at 5 wpf. The framed regions are enlarged to show details on the right with single channel images shown in grayscale. Arrows indicate representative mCherry^+^. Scale bar, 50 μm. **d,** Experimental design for (e, f, g). **e, f,** Fate mapping results showing HCR staining of *col18a1a* (e, red), *pdgfrb* (e, blue), and *vegfaa* (f, red) in lineage-traced EGFP^+^ cells (green). Framed regions are enlarged to show representative colocalizations (arrows). Scale bar, 50 μm (e) or 20 μm (f). **g,** Quantification of cell fates as shown in (e). **h,** Experimental design for (i-j). **i,** Fate mapping results showing HCR staining of *col18a1a* (red) and *pdgfrb* (blue) in lineage-traced EGFP^+^ cells (green). Framed regions are enlarged to show representative colocalizations (arrows and arrowhead) with single-channel images shown in grayscale. Scale bar, 25 mm. **j**, Quantification of EGFP^+^ cells as shown in (i).

### *col18a1a*^+^ mesenchymal cells are epicardial-derived perivascular cells distinct from mural cells

To better understand the *col18a1a*^+^ progeny and their relationship with the coronary vasculature, we performed HCR staining of endogenous *col18a1a* in the adult heart. As shown in Fig. 4a, *col18a1a*^+^ cells are adjacent to coronary vessels, suggesting a perivascular cell identity. To further label these cells, we generated a *col18a1a:EGFP* BAC reporter and characterized its expression. In the adult heart, *col18a1a*:EGFP expression was observed across the entire ventricle in *tcf21*:H2A/Z-mCherry (or *tcf21*:H2R)^+^ cells, suggesting an epicardial origin (Fig. 4b). We crossed the line with the arterial reporter *flt1:TdTomato* and observed that *col18a1a*^+^ cells wrap around arteries (Fig 4c, arrows). *col18a1a*^+^ cells were also found contacting vessels not visualized by *flt1*:TdTomato, likely the coronary veins or capillaries (Fig 4c, arrowheads). These observations validate that the EGFP reporter faithfully recapitulates endogenous expression. To further confirm the identity of *col18a1a*^+^ cells, we performed HCR staining of *pdgfrb*, the mural cell marker. As shown in Fig. 4d, *col18a1a*^+^ cells exhibit minimal *pdgfrb* expression compared with *pdgfrb*^high^ mural cells. Our single-cell transcriptomic data indicated that *col18a1a*^+^ cells (clusters 0, 1, and 4) have no or minimal expression for several mural cell (pericyte and VSMC, cluster 6) markers, such as myosin heavy chain 11a (*myh11a*), tropomyosin 1 (*tpm1*), regulator of G protein signaling 5a (*rgs5a*), *notch3*, and NDUFA4 mitochondrial complex associated like 2a (*ndufa4l2a*)^40,41^ (Fig. 4e and Supplementary Fig. S1). These finding suggest that *col18a1a* is a marker for a perivascular cell fate that is distinct from pericytes and VSMCs and largely originates from epicardial progenitors transitioning through *scxa* (epicardial-derived perivascular mesenchymal cells or Epi-PMCs). We next assessed the expression dynamics of *col18a1a* during coronary angiogenesis in juvenile hearts. We observed that *col18a1a*:EGFP expression emerged near the AVC around 6 wpf, a stage when coronary vessels had already expanded across the ventricular surface but mirroring their earlier development from AVC to ventricle surface (Fig. 4f). Given that the COL18A1-derived endostatin peptide is known to stabilize newly formed endothelial tubes and inhibit angiogenesis^36–38^, *col18a1a* expression likely reflects vascular maturation, paralleling the spatiotemporal wave of vessel development that begins near the AVC^30,42,43^.

**Figure 4.**
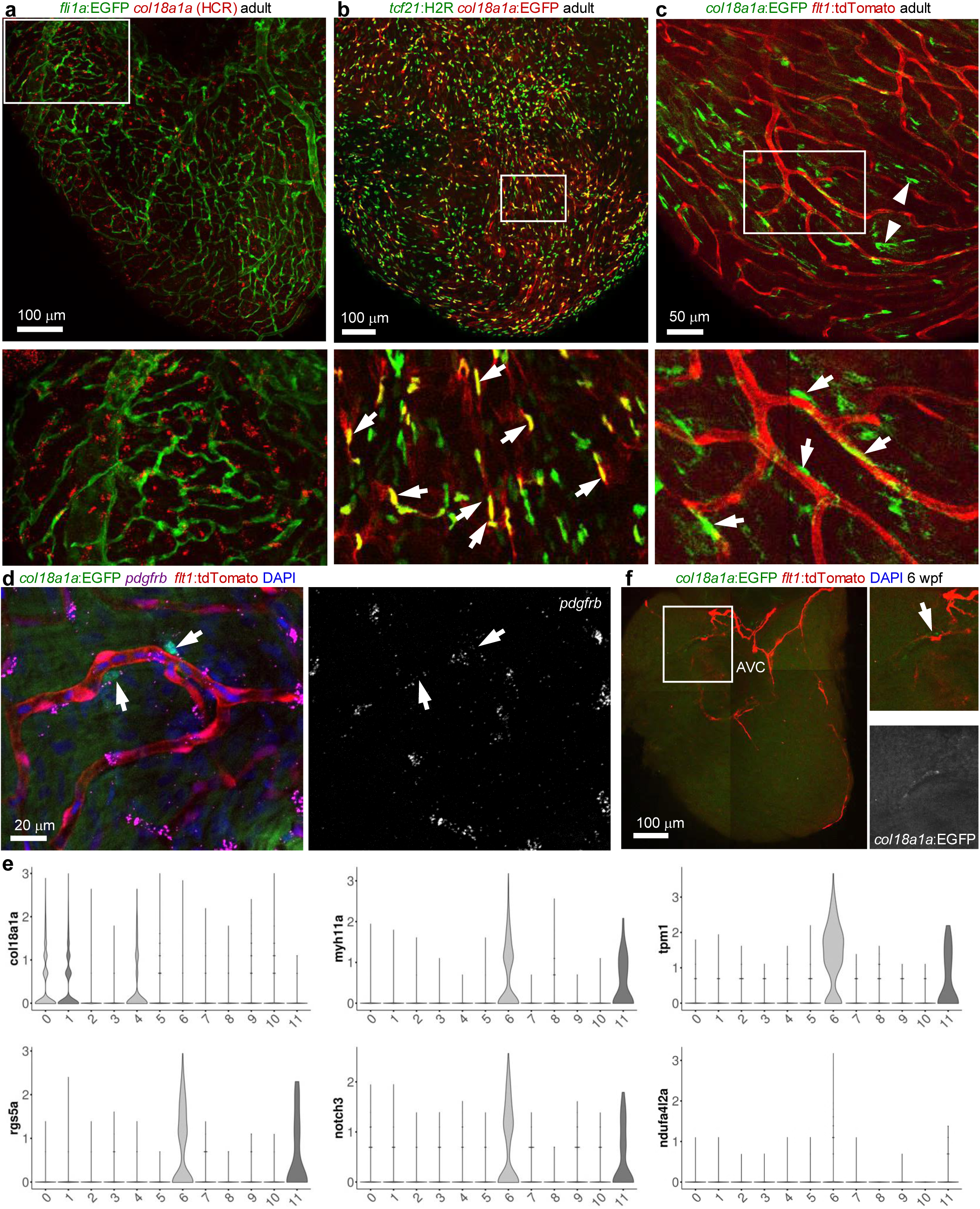
*col18a1a*^+^ cells are perivascular cells distinct from pericytes and vascular smooth muscle cells. **a**, A whole-mount image of the ventricular surface showing HCR staining of *col18a1a* (red) and *fli1a*:EGFP (green) expression in an adult heart. The framed region is enlarged to show details at the bottom. Scale bar, 100 μm. **b**, A whole-mount image of the ventricular surface showing expressions of *tcf21*:H2R (green) and *col18a1a*:EGFP (red) in an adult heart. The framed region is enlarged to show details at the bottom. Arrows indicate representative H2R^+^ EGFP^+^ cells. Scale bar, 100 μm. **c**, A whole-mount image of the ventricular surface showing expressions of *col18a1a*:EGFP (green) and *flt1*:tdTomato (red) in an adult heart. The framed region is enlarged to show at the bottom. Arrows indicate representative EGFP^+^ cells in adjacent to coronary vessels. Scale bar, 50 μm. **d**, A whole-mount image of the ventricular surface showing HCR staining signals of *pdgfrb* in magenta. Expressions of *col18a1a*:EGFP and *flt1*:tdTomato are shown in green and red, respectively. DAPI staining is shown in blue. A single-channel image of *pdgfrb* signals is shown in grayscale on the right. Arrows indicate EGFP^+^*pdgfrb*^low^ cells. Scale bar, 20 μm. **e,** Violin plots displaying the expression patterns (log1p) of selected marker genes across cell clusters. **f,** A whole-mount image of the ventricular surface showing expressions of *col18a1a*:EGFP (green) and *flt1*:tdTomato (red) in a 6 wpf heart. The framed region is enlarged to show details on the right with the single-channel image shown in grayscale. Arrows indicate representative EGFP^+^ cells in adjacent to the AVC (dashed circle). Scale bar, 100 μm.

### Loss of *scxa* affects coronary angiogenesis but not cardiomyocyte proliferation

To determine the role of *scxa* in coronary angiogenesis we analyzed the coronary vasculature using the *fli1a:EGFP* reporter in a global mutant *scxa^kg170^* (Ref. ^44^). Quantification of vascular area in adult ventricles revealed a trend of increased vascular density in *scxa^-/-^* mutants compared to wild-type siblings (Fig. 5a, b). However, the number of *col18a1a*^+^ cells seems comparable in the mutant and wild-type siblings (Supplementary Fig. S2a, b). We further examined whether *scxa* deletion influences CM proliferation during heart regeneration by subjecting *scxa^-/-^* and wild-type sibling fish to ventricular amputation. However, CM proliferation indices were comparable between the two groups at 7 dpa (Fig. 5c, d). To address potential genetic compensation by *scxb*, whose expression was detected in the zebrafish heart (Supplementary Fig. S2c), and to assess the collective role of *scleraxis* genes in coronary angiogenesis, we employed *scxa^kg170^scxb^kg107^* double mutants^44^. These mutants displayed severe musculoskeletal defects and high mortality, typically dying before 3 wpf, likely due to mandibular cartilage defects impairing feeding^44^. However, by isolating and providing excessive feeding, we successfully raised <5% of the double mutants to 3 mpf, although they are substantially smaller in body and heart sizes than their wild type siblings (Fig. 5e). Notably, these viable mutants lacked coronary vessels (Fig. 5f). While the absence of coronary vessels in the double mutants suggests that *scleraxis* genes may contribute to angiogenesis, this phenotype may in part arise from indirect developmental defects. Combined with the low survival rate, these limitations prevented further mechanistic analysis of the double mutants.

**Figure 5.**
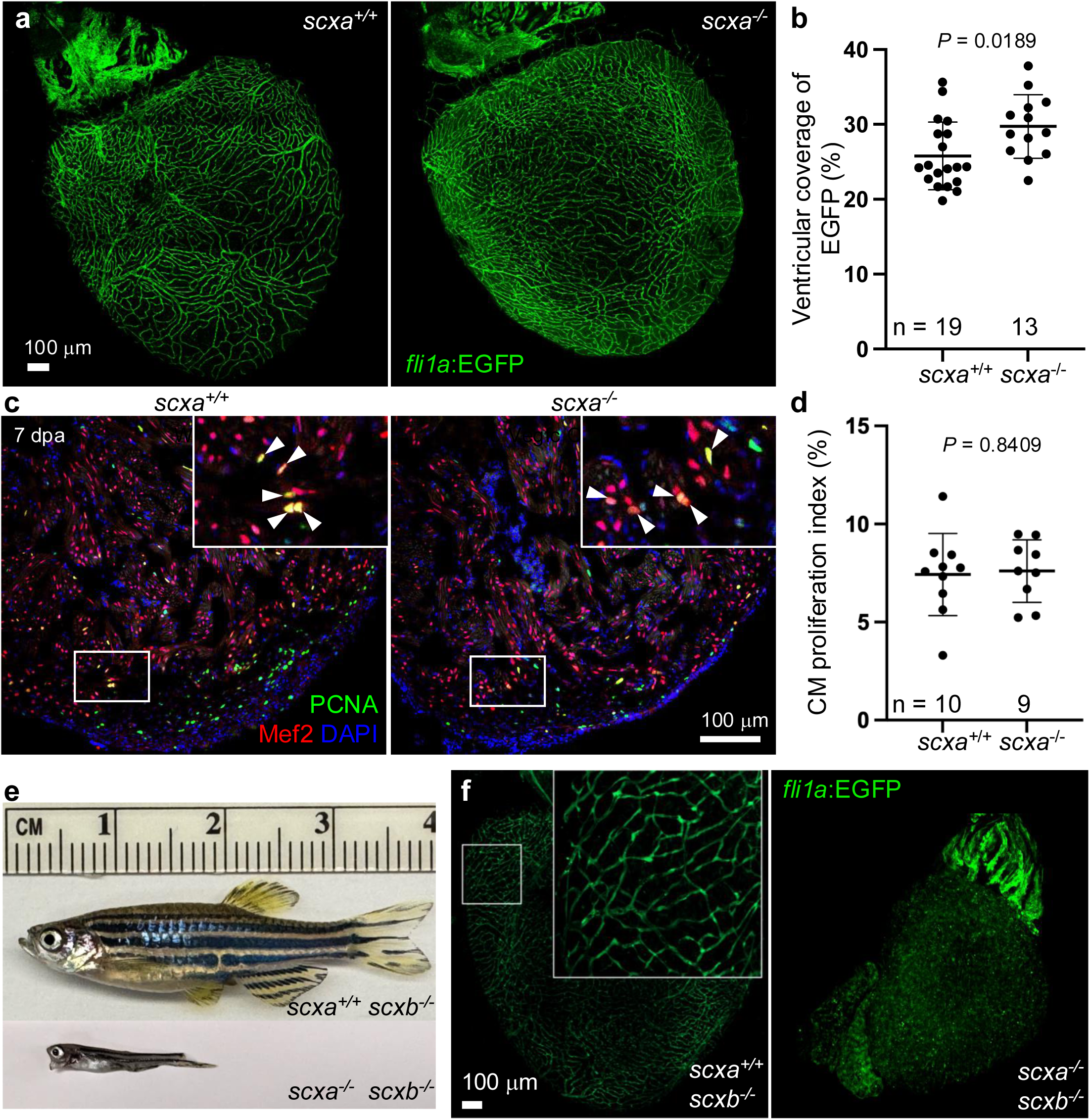
Loss of *scxa* affects coronary angiogenesis but not cardiomyocyte proliferation. **a**, Whole-mount images showing ventricular coverage of *fli1a*:EGFP vessels (green) in WT (left) and *scxa*^-/-^ hearts (right). Scale bar, 100 μm. **b,** Quantification of ventricular vessel coverage as shown in (a). Student’s *t*-test. **c,** Cryosection images showing antibody staining for PCNA (green) and Mef2 (magenta). The framed regions are enlarged and displayed at the top. Dashed lines denote injury sites. The arrowheads indicate representative PCNA^+^Mef2^+^ cells. Scale bar, 100 μm. **d,** Quantification of the CM proliferation index. Student’s *t*-test. **e,** *scxa*^-/-^*scxb*^-/-^ fish exhibit developmental defects. **f**, Whole-mount images showing ventricular coverage of *fli1a*:EGFP vessels (green) in WT (left) and *scxa*^-/-^*scxb*^-/-^ (hearts). Scale bar, 100 μm.

### Hypoxia induces *scxa* expression

To identify factors regulating *scxa* expression, we performed CUT&Tag analysis of histone H3K4me1 modifications in aEPCs. We isolated aEPCs at 3 dpa using *tcf21:nucEGFP* and *ptx3a:mScarltet-NTR* reporter for EGFP^+^mScarlet^+^ cells (Fig. 6a). EGFP^+^ cells from uninjured hearts were used as a control for quiescent epicardial cells. We combined the new CUT&Tag data with our published RNA-seq, ATAC-seq, and Chip-seq results^45^. As shown in Fig. 6b, we observed 4 putative promoter and/or enhancer regions upstream of the *scxa* gene that gained accessibility and enhancer markers (H3K4me1 and H3K27Ac) after heart injury. Motif analysis using JASPAR^46^ predicted multiple putative HIF1A (Hypoxia-Inducible Factor 1 Alpha)-binding sites within these regions (Fig. 6c; profile score > 4), implying hypoxia and Hif signaling as regulators of *scxa* expression. To test the impact of excessive hypoxia on *scxa* expression, we used phenylhydrazine (PHZ) to induce acute hemolysis and hypoxia^47–49^. After treatment with PHZ (1.5 mg/ml for 20 min, every other day for seven days), we observed a striking re-expression of *scxa*:mCherry expression in the entire ventricular surface (Fig. 6d). Importantly, the ectopic *scxa* expression is enriched adjacent to coronary vessels (Fig. 6d insert). To evaluate whether this effect is reproducible in juvenile heart development, we applied PHZ treatment to juveniles for one week from 7 wpf and assessed *scxa* expression at 8 wpf. As expected, PHZ treatment significantly increased *scxa* expression in juveniles (Fig. 6f, g). To further test the role of HIF-mediated hypoxic signaling, we treated juveniles with 0.1 mM Dimethyloxalylglycine (DMOG) from 6 to 8 wpf, which stabilizes Hif factors and inducec hypoxic responses^50,51^. Compared with vehicle-treated controls, DMOG treatment resulted in increased *scxa* expression (Fig. 6h, i). Altogether, our results suggest that hypoxia and Hif signalinginduces *scxa* expression, thereby promoting aEPC differentiation and contributing Epi-PMCs to the coronary network.

**Figure 6.**
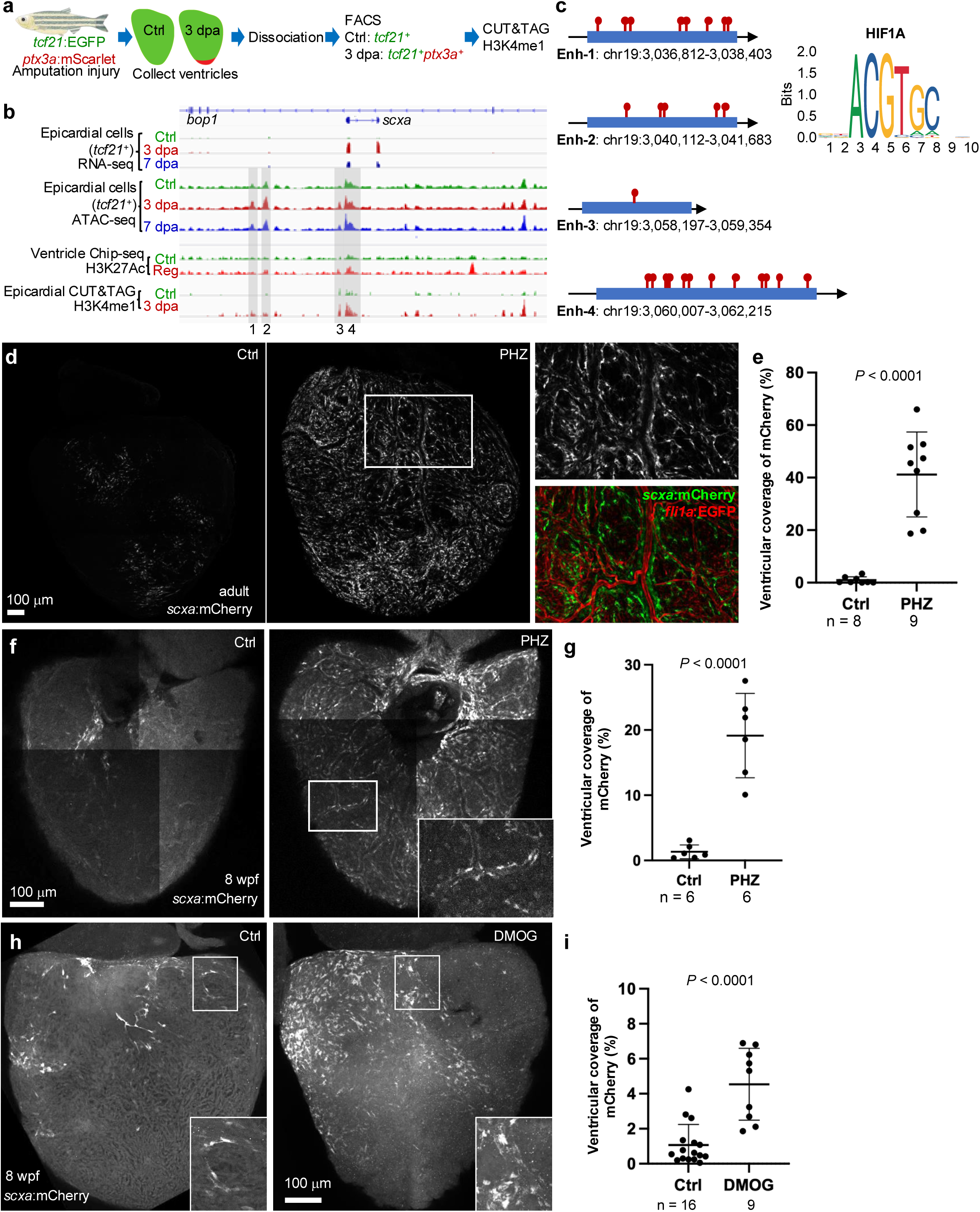
Hypoxia induces *scxa* expression in adults. **a**, Schematic for experiment design. Heart resection injuries were performed in *tcf21:nucEGFP; ptx3a:mScartlet* animals. Ventricles were collected at 3 dpa together with those of uninjured siblings (Ctrl). Ventricles were dissociated, and EGFP^+^ or EGFP^+^ mScarlet^+^ epicardial cells were isolated by FACS for the CUT&TAG assay. **b**, A browser track of the genomic region comprising gene *scxa* showing the transcripts, chromatin accessibility profiles, and CUT&TAG marks of H3K4me1 in the epicardium across replicates from Ctrl, 3 dpa, and 7 dpa samples. The whole-ventricle H3K27Ac profile of the uninjured (Ctrl) and regenerating (Reg) hearts is shown. Gray boxes indicate putative enhancer regions with increased accessibility and enhancer markers during regeneration. **c**, Putative HIF1A binding sites in the 4 enhancer (Enh) regions shown in (b). **d**, Whole-mount images showing *scxa*:mCherry expressions without (Ctrl, left) or with PHZ treatment (middle). The framed region is enlarged to show detail on the top right with the coronary vessel marker *fli1a*:EGFP shown in red in the overlayed image (bottom right). Scale bar, 100 μm. **e**, Quantification of ventricular coverage of *scxa*:mCherry signals as shown in (d). Student’s *t*-test. **f**, Whole-mount images showing *scxa*:mCherry expressions without (Ctrl, left) or with PHZ treatment (right). The framed region is enlarged to show detail on the bottom right Scale bar, 100 μm. **g**, Quantification of ventricular coverage of *scxa*:mCherry signals as shown in (f). Student’s *t*-test. **h,** Whole-mount images showing *scxa*:mCherry expressions with vehicle (Ctrl, left) or DMOG (right) treatment at 8 wpf after a 2-week treatment. The framed regions are enlarged in the bottom right for details. Scale bar, 100 mm. **i,** Quantification of ventricular coverage of *scxa*:mCherry signals at 8 wpf. Student’s *t*-test.

## DISCUSSION

The epicardium is recognized as a dynamic and multipotent contributor to heart development, repair, and regeneration. In this study, we identify Scxa as a critical transitional regulator within the epicardial lineage that governs differentiation toward a previously unexplored *col18a1a*^+^ perivascular population Epi-PMCs (Fig. 7e). These cells are molecularly distinct from canonical mural cell types and are characterized by a unique combination of extracellular matrix components and angiogenic regulators. Our findings expand the spectrum of epicardial-derived fates and shed light on how transcriptional and environmental inputs are integrated during coronary vascular morphogenesis.

**Figure 7.**
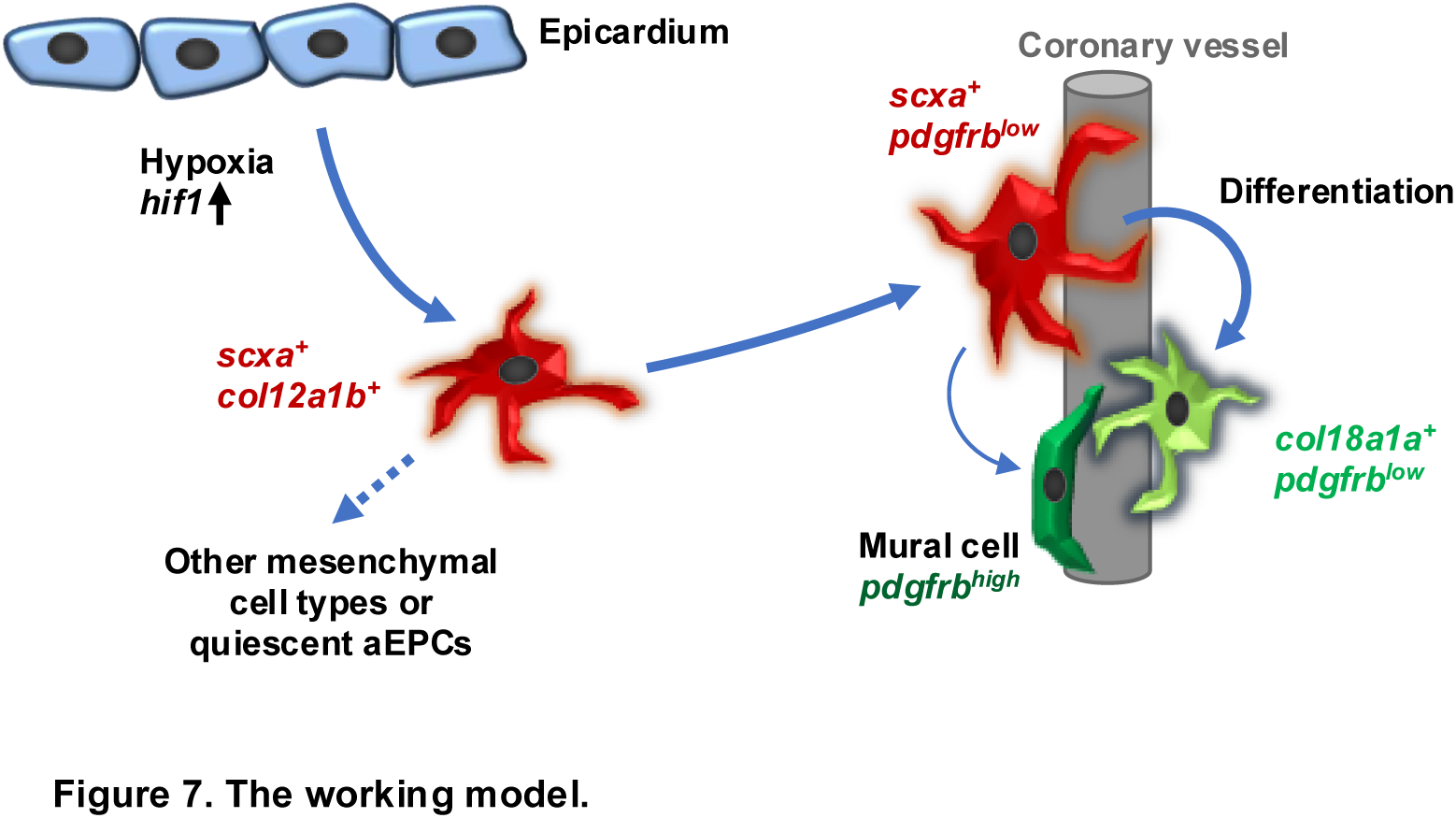
The working model.

Although Scx is historically associated with tendon development, emerging evidence links it to cardiac processes. In mammals, Scx has been linked to pro-epicardial organ development, embryonic epicardial cells, cardiac fibroblast formation, and valve morphogenesis^21–26^. Its prolonged activation post-injury promotes a fibrotic myofibroblast phenotype through upregulation of fibrillar collagen (Col1a2)^26^. Interestingly, loss of Scx in mice results in hypocellular, underdeveloped hearts, indicating a broader role in promoting mesenchymal cell differentiation^26^. These seemingly opposing outcomes underscore a key principle: Scx functions are highly context- and time-dependent, and its cellular effects are shaped by local signaling environments and lineage states. In the zebrafish heart, we find that *scxa* expression is transient and tightly linked to a hypoxic microenvironment. This limited temporal window may allow epicardial cells to flexibly adopt supportive roles in angiogenesis. In contrast, the adult mammalian epicardium likely lacks this flexibility and typically responds to injury by generating fibrotic tissue rather than functional vasculature. Understanding how *scxa* is temporally gated and environmentally regulated could provide valuable insights into the failure of regenerative programs in the adult human heart.

Our results also underscore a growing appreciation for the functional diversity of perivascular cells, which extend beyond canonical pericytes and vascular smooth muscle cells. A previous study by Katz et al. using Scx lineage tracing suggested an endothelial cell fate for Scx^+^ proepicardial cells^23^. However, although we are yet to define the fate of the remaining ∼14% of *scxa*-linage (Fig. 3g), our study suggests a primarily perivascular rather than endothelial cell fate of *scxa*^+^ epicardial progenitors. Whether these *col18a1a*^-^*pdgfrb*^-/low^ cells represent a quiescent progenitor population that can be further activated upon heart injury remains an important question for future investigation. Previous studies in zebrafish have described perivascular fibroblast populations in the trunk vasculature that exhibit low *pdgfrb* expression and contribute to vessel stability^52^. These cells were shown to support vessel integrity and, in some cases, differentiate into pericytes. The *col18a1a*^+^ Epi-PMCs identified here resemble these perivascular fibroblasts in expression profile and morphology but likely carry distinct functions suited to the complexity of the coronary vasculature. Notably, Col18a1a encodes collagen XVIII, a basement membrane component that can be proteolytically cleaved to yield endostatin, an anti-angiogenic peptide that stabilizes endothelial tubes^36–38^. Simultaneously, Epi-PMCs discovered in our study also express *vegfaa*, a potent pro-angiogenic factor. This duality suggests that *col18a1a*^+^ cells may serve a regulatory role, balancing vessel sprouting with maturation. In support of this, Col18a1 knockout mice show increased vascular density in the retina^53^, implicating this matrix protein in vascular remodeling. Our observed late onset of *col18a1a* expression during coronary angiogenesis in zebrafish further supports its role in the maturation phase rather than the initiation of vessel growth. Together, these findings highlight a more active and responsive role for the epicardial lineage in shaping vascular architecture through intermediate cell types with specialized signaling outputs.

The increased coronary vessel coverage observed in *scxa* single mutants suggests that Scxa may act, at least in part, as a negative regulator of angiogenesis in our experimental setting. This finding is consistent with the proposed role of Col18a1a-derived endostatin, an anti-angiogenic peptide produced by *scxa*^+^ perivascular cells. However, the number of *col18a1a*^+^ cells remained unchanged in *scxa* mutants. Possible explanations include decreased *col18a1a* expression, unopposed pro-angiogenic signals such as Vegfaa within these cells, compensatory effects from *scxb*, and reduced enzymatic processing of endostatin. Notably, *scxa*^+^ cells also express *vegfaa* before differentiating into *col18a1a*^+^ cells (Fig. 3a). While further validation is needed, these results support a model in which Scxa modulates angiogenesis by temporally balancing vessel-promoting and vessel-stabilizing signals within the epicardial-derived cell population. Notably, whereas *scxa* mutants exhibited increased vessel coverage, coronary vessels were entirely absent in *scxa*/*scxb* double mutants, indicating a redundant but essential role for *scleraxis* genes in coronary vessel formation. Although systemic developmental defects may contribute to the double mutant phenotype, the contrasting outcomes may highlight the context-specific functions of Scx transcription factors, ranging from fine-tuning vessel growth to enabling vascular initiation.

One of the intriguing aspects of this study is the connection between hypoxia and *scxa* induction. Our epigenomic analyses identified multiple HIF1A motifs in injury-responsive putative regulatory regions near *scxa*, and systemic induction of hypoxia dramatically elevated *scxa* expression in epicardial cells. These findings align with previous reports showing that hypoxia induces epicardial cell differentiation and vascular precursor cell invasion into the myocardium^54–56^. It was recently found that stabilization of hypoxic signaling in the epicardium promotes regenerative outcomes in mammalian models of myocardial infarction^57^. By linking HIF signaling to a defined transcriptional program and cell fate, our data provide a mechanistic bridge between hypoxia and epicardial plasticity.

In conclusion, this study identifies a hypoxia-responsive Scxa-Col18a1a axis that promotes the differentiation of epicardial progenitors into a perivascular mesenchymal population involved in coronary vessel formation. Beyond revealing new lineage complexity within the epicardium, our findings highlight Scxa as a potential molecular handle to modulate regenerative responses in the injured heart. The transient expression of *scxa* in zebrafish may be a key feature enabling regenerative plasticity. Understanding and reconstituting this regulatory program could be central to restoring vascular integrity in ischemic heart disease and improving regenerative therapies.

## METHODS

### Animal Maintenance and Procedures

All animal protocols were approved by the Institutional Animal Care and Use Committee (IACUC) at Weill Cornell Medical College. Adult zebrafish (aged 3-12 months) of the Ekkwill (EK) and EK/AB strains were maintained as previously described^58,59^. Animals of both sexes were used for experiments. Water temperature was maintained at 28°C, and fish were kept on a 14/10 light/dark cycle at a density of 5-10 fish per liter. Heart amputation injury was performed as previously described^60^. For tamoxifen treatments, juvenile and adult fish were placed in a mating tank of aquarium water containing 5 μM 4-Hydroxytamoxifen (4HT, Sigma-Aldrich). Embryos were treated with 5 μM 4HT in dishes. Fish were maintained for 16 h, rinsed with fresh aquarium water, and returned to the recirculating aquatic system. For phenylhydrazine (PHZ, Sigma-Aldrich) treatment, adult fish were placed in a mating tank of aquarium water containing 1.5 μg/ml PHZ for 20 min daily. Fish were rinsed with fresh aquarium water and returned to the recirculating aquatic system. For all reporter lines, fish were analyzed as hemizygotes. The following published lines were used: *Tg(tcf21:nucEGFP)^pd41^* (Ref. ^32^), *Tg(tcf21:H2A-mCherry)^pd252^* (Ref. ^61^), *Tg(tcf21:CreER^t2^)^pd42^* (Ref. ^32^), *Tg(pdgfrb:EGFP)^ncv22^* (Ref. ^33^), *Tg(col12a1b:EGFP)^wcm108^* (Ref. ^15^), *Tg(ptx3a:mScarlet-P2A-NTR)^wcm107^*(Ref. ^15^), *TgBAC(ubb:loxP-TagBFP-STOP-loxP-EGFP)^vcc18^*(Ref. ^31^), *Tg(scxa:mCherry)^fb301^*(Ref. ^27^), *Tg(scxa:CreER^t2^)^xxx^*(Ref. ^39^), *scxa^kg170^*(Ref. ^44^), and *scxb^kg107^*(Ref. ^44^). Newly generated lines are described below. The generation of new lines is described below. All reporters were analyzed as hemizygotes.

### Generation of *Tg(col18a1a:EGFP)* zebrafish

To make a BAC construct, the translational start codon of *tfa* in the BAC clone xxx was replaced with the *EGFP-polyA* cassette by Red/ET recombineering technology (Gene Bridges)^62^. The 5’ and 3’ homologous arms for recombination were a 50-base pair (bp) fragment upstream and downstream of the start codon and were included in PCR primers to flank the *EGFP-polyA* cassette. The primer sequences are: Forward, 5’-CAACAACAATCCCATGTAGCTATCGGGACGGAGCAGTTTCACGAACTGAAATGGTGAGCAAGGGCGAGGAGCTG-3’; Reverse, 5’- CTTAAAATATATACTCCAAAACTCATGTATCCTCATGTATACCTTCTTGCGCCCTTGAAGTTCCTATTCTCTAG-3’. The final BAC was purified with Nucleobond BAC 100 kit (Clontech) and co-injected with PI-Sce I into one-cell-stage zebrafish embryos. Stable transgenic lines with bright fluorescence were selected. The full name of this line is *Tg(col18a1a:EGFP)^wcm113^*.

### CUT&Tag assay and data analysis

CUT&TAG assay targeting H3K4me1 was performed using the EpiCypher CUTANA™ CUT&Tag Kit following the manual and a published protocol ^63^. Briefly, heart amputation surgery was performed on adult fish (6 months of age) carrying *tcf21:nucEGFP* and *ptx3a:mScarlet-P2A-NTR* reporters. Hearts were isolated at 3 dpa together with the uninjured control (2 biological replicates each) on ice and washed several times to remove blood cells. Ventricles were digested in an Eppendorf tube with 0.5 ml HBSS plus 0.13 U/ml Liberase DH (Roche) at 37°C, while stirring gently with a Spinbar® magnetic stirring bar (Bel-Art Products). Supernatants were collected every 5 min and neutralized with sheep serum. Dissociated cells were spun down and re-suspended in DMEM plus 10% fetal bovine serum (FBS) medium and sorted using a Sony MA900 sorter for EGFP^+^ cells (uninjured) and EGFP^+^mScarlet^+^ (3 dpa). Nuclei preparation and CUT&TAG library preparation using rabbit anti-H3K4me1 antibody (Abcam, ab8895) were performed using the EpiCypher CUTANA™ CUT&Tag Kit following the manual. Two replicates with 5,000-12,000 nuclei for each sample were used. Sequencing was performed using an Illumina NextSeq2000 with 11 - 44 million 50-nt pair-end reads obtained for each library. CUT&TAG data were processed and analyzed following a published pipeline^64,65^. Briefly, reads were trimmed for adaptors using Trimmomatic (v0.39) before aligning to the DanRer10 genome using Bowtie2 (v2.5.3) with parameters for 10-700 bp length. Peaks were called using SEACR (v1.3) with p-value < 0.01.

### Histology and Microscopy

Freshly isolated hearts were fixed overnight at 4°C in 4% paraformaldehyde (PFA). Fixed hearts were either mounted with Fluoromount G (Southern Biotechnology, cat#0100-01) between two coverslips to visualize both ventricular surfaces or processed for cryosectioning at a thickness of 10 μm. Hybridization Chain Reaction (HCR) staining was performed on whole-mount hearts and cryosections following established protocols^66^. HCR probes targeting *col18a1a* and *pdgfrb* were obtained from Molecular Instruments Inc. Immunostaining of whole-mount hearts and cryosections was carried out as previously described^61,67^. The primary antibodies used in this study included mouse anti-GFP monoclonal antibody (3E6) (Thermo Fisher, A11120), rabbit anti-dsRed/mCherry (1:200, Takara, 632496), and chick anti-GFP (Aves Labs, GFP-1020). Secondary antibodies (Thermo Fisher) included Alexa Fluor 488 goat anti-rabbit, goat anti-mouse, goat anti-chick, Alexa Fluor 546 goat anti-rabbit, goat anti-mouse, and Alexa Fluor 633 goat anti-mouse. Fluorescent images of whole-mount and cryosectioned heart samples were acquired using a Zeiss LSM 800 confocal microscope.

### RT-PCR

Five hearts per sample were dissected, washed in sterile PBS, collected in 500 μl QIAzol (Qiagen), and mixed with 100 μl of 0.5 mm RNase-free stainless-steel beads. Samples were homogenized using a Next Advance Bullet Blender for 5 minutes and subsequently stored at -80 °C. Total RNA was extracted using the RNeasy Mini Kit (Qiagen) according to the manufacturer protocol. cDNA synthesis was performed with the SuperScript III First-Strand Synthesis Kit (Invitrogen) following the manual. PCR amplification of target genes was conducted using the DreamTaq PCR system under the following conditions: 30 cycles with an annealing temperature of 59 °C. Primer sequences were as follows:

*scxa*-forward: 5′-GTCAACGCAATGCTGCCAAT-3′;

*scxa*-reverse: 5′-TCAGCAGGCTCTGTAGGGAT-3′;

*scxb*-forward: 5′-GCGAGGGTTCTGGATCTGAG-3′;

*scxb*-reverse: 5′-ACTGTTCGTTCGTTCTCGCT-3′;

*gapdh*-forward: 5′-GATGGTCATGCAATCACAGTCTA-3′;

*gapdh*-reverse: 5′-ATCATACTTGGCAGGTTTCTCAA-3′;

*rpl13a*-forward: 5′-TCTGGAGGACTGTAAGAGGTATGC-3′;

*rpl13a*-reverse: 5′-AGACGCACAATCTTGAGAGCA-3′.

### Data collection and statistics

For each treatment, clutchmates were randomly assigned to different experimental groups. All experiments included a minimum of two biological replicates. Sample sizes were determined based on prior studies and the nature of the experiments, with details provided in each figure legend. Statistical results are presented as the mean ± standard deviation (s.d.). Information on sample sizes, statistical tests, and P-values can be found in the figures or their corresponding legends. Two-tailed Student’s *t*-tests were used for comparisons when assumptions of normality and equal variance were satisfied.

### Data availability

The CUT&Tag datasets generated in this study have been deposited at NCBI’s Gene Expression Omnibus (GEO) under accession numbers GSE299367. The previously published RNA-seq and ATAC-seq datasets of epicardial cells are available under access numbers GSE89444. Ventricle Chip-seq (H3K27Ac) dataset was downloaded from GEO under accession number GSE75894.

## ACKNOWLEDGMENTS

We thank A. Afolalu, C. Shapiro, S. Hosten, and C. Quaies for fish care, Dr. Jenna Galloway for *scxa:mCherry* and *scxa:CreER^t2^* lines, Dr. Yaniv Hinits for *scxa^kg170^* and *scxb^kg107^* mutants, Junsu Kang for comments on the manuscript. This work was supported by the New York State Stem Cell Science program (NYSTEM) predoctoral training grant position to B.P., The Louis and Rachel Rudin Foundation fellowship to Y.X., a Research Fellowship award from the Belgian American Educational Foundation to I.V.W. American Heart Association (AHA) Career Development Award (AHA941434) and National Institutes of Health (NIH) grant R01NS126209 to M.R.H., AHA Transformational Project Award (23TPA1077772) and NIH grants (R01HL155607, R01HL166518) to J.C.

## AUTHOR CONTRIBUTIONS

Conceptualization: J.C.

Methodology: B.P., Y.X., P.Z., D.B., and J.C.

Investigation: B.P., Y.X., J.Y., M.Q., A.G.C.Y., M.N., I.V.W., and A.Y.

Resources: K.K., M.R.H. and J.C.

Writing and editing: B.P., T. E., M.R.H., and J.C.

Funding Acquisition: M.R.H. and J.C.

---

**Supplementary Figure S1.**
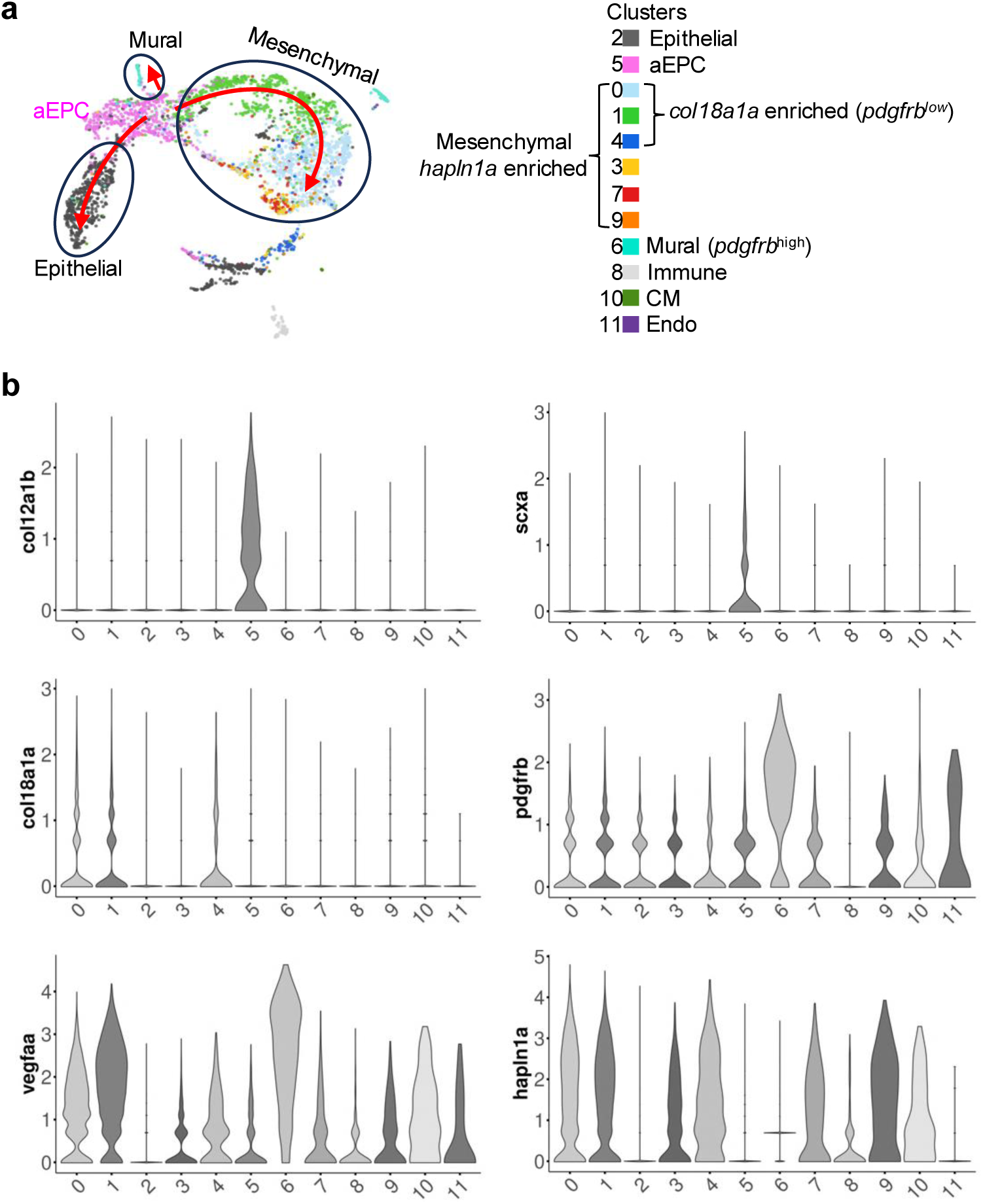
Gene expression analysis from single-cell RNA-seq data. **a,** Pseudotime trajectories inferred using Monocle 3, visualized on a UMAP plot, highlight cell differentiation paths and cell types. **b**, Violin plots displaying the expression patterns (log1p) of selected marker genes across cell clusters.

**Supplementary Figure S2.**
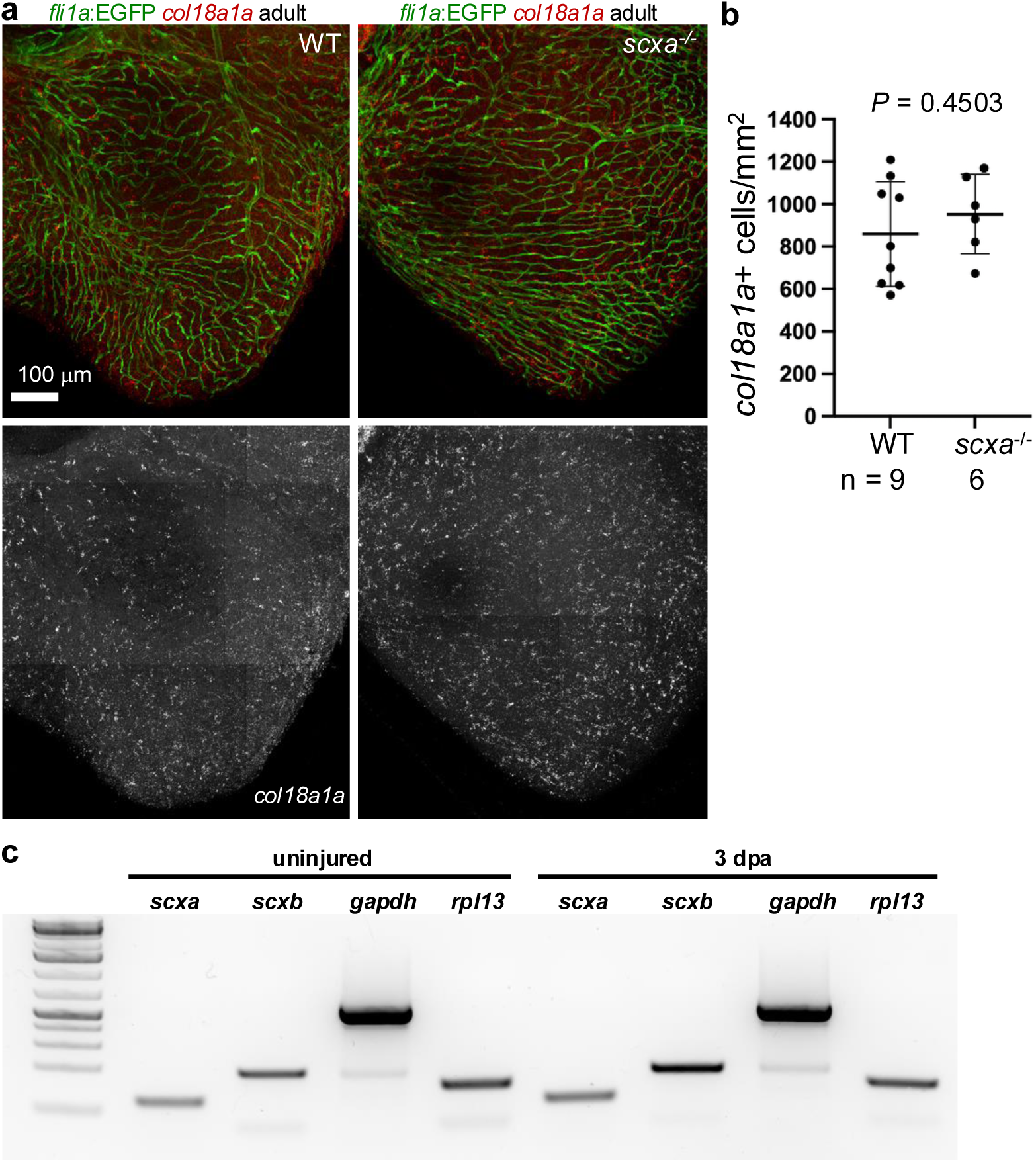
The *scxa* mutant. **a**, Whole-mount images of ventricles showing *fli1a*:EGFP vessels (green) and HCR staining of *col18a1a* (red) in WT (left) and *scxa*^-/-^ hearts (right). Scale bar, 100 μm. **b,** Quantification of *col18a1a*^+^ cell density as shown in (a). Student’s *t*-test. **c**, RT-PCR results of selected genes using wild-type adult hearts collected at 3 dpa together with the uninjured control. Five hearts were used for each group.

